# Reconstruction of the gastropod ancestral karyotype reveals lineage-specific chromosomal evolution and adaptations during habitat transitions

**DOI:** 10.64898/2026.09.02.748897

**Authors:** Zhaoyan Zhong, Yufei Zhou, Hui Wang, Chong Chen, Xu Liu, Jin Sun

## Abstract

Gastropoda is the most diverse class of molluscs, with spectacular adaptive radiations across shallow to deep marine, freshwater, and terrestrial environments, yet the role of chromosomal restructuring behind these habitat transitions remain largely unexplored. This is in part due to most chromosomal genomes available are in the subclasses Caenogastropoda and Heterobranchia, with few representatives from the other four subclasses. Here, we assembled chromosome-level genomes for five representatives from those four subclasses: *Vittina coromandeliana* (Neritimorpha), *Shinkailepas gigas* (Neritimorpha), *Turbo cornutus* (Vetigastropoda), *Neomphalus fretterae* (Neomphaliones), and *Cellana toreuma* (Patellogastropoda), and reconstructed the ancestral gastropod karyotype consisting of 20 linkage groups (GLGs) as well as the ancestral karyotype of each subclass. Integrating macrosynteny and phylogenomic analyses across all six subclasses, we show that different lineages have adopted fundamentally different chromosomal strategies to colonize novel habitats: freshwater caenogastropods genomes are characterized by extensive end-to-end fusions, terrestrial heterobranchs by whole-genome duplications, freshwater neritimorphs by lineage-specific fissions plus fusion-with-mixing, and deep-sea hydrothermal vent vetigastropods by widespread chromosomal restructuring, whereas hydrothermal vent neomphaliones and caenogastropods retain remarkably conserved ancestral karyotypes. Rearrangement breakpoints preferentially reside within topologically associating domains (TADs) rather than at boundaries, preserving higher-order chromatin architecture. Functionally, convergent enrichment of the mucin-type O-glycan biosynthesis pathway in both freshwater and hydrothermal vent lineages is accompanied by lineage-specific modifications, enhanced immune modulation in freshwater versus reinforced physicochemical shielding in the deep sea, reflecting divergent adaptive applications of this pathway. Our findings demonstrate that habitat transitions in gastropods are facilitated not by a uniform genomic mechanism, but by a diverse repertoire of lineage-specific chromosomal strategies, highlighting the extraordinary genomic flexibility that underpins the remarkable gastropod diversity seen today.

## Introduction

Rates of speciation and diversification vary substantially among animal clades, and also through time; adaptive radiations can lead to increased speciation rates and ecological diversification (Schluter 2000; Losos 2010). Environmental shifts, mass-extinction aftermaths, and the colonization of new habitat types with a broad range of available niches can all provide ecological opportunities for rapid radiation (Vermeij 1977; Raup and Sepkoski 1982; Marshall 2006). During the Ediacaran-Cambrian radiation, pulses of shallow ocean oxygenation events may have expanded habitable space and, together with gene regulation and ecological interactions contributed to an explosive diversification (Erwin et al. 2011; Zhang and Shu 2014; Bowyer et al. 2024). The Cretaceous-Paleogene extinction event eliminated many existing lineages and was followed by rapid radiation and lineage splitting in surviving groups such as modern birds and placental mammals (Jarvis et al. 2014; Hull 2015; Foley et al. 2023). Adaptive radiation coupled with colonization of drastically different ecosystems, such as marine to freshwater transitions, or island-like habitats can also lead to rapid adaptive radiation (Lee and Bell 1999; Cerca et al. 2022), sometimes coupled with drastic changes in the genomic architecture (Lewin et al. 2024; Vargas-Chávez et al. 2025).

Mollusca is the second largest animal phylum encompassing eight classes, with Gastropoda containing familiar animals like limpets, snails, and slugs being the most species-rich (Chen et al. 2025b). Currently, Gastropoda comprises six subclasses: Patellogastropoda, Vetigastropoda, Neomphaliones, Caenogastropoda, Neritimorpha, and Heterobranchia (Bouchet et al. 2017); the number of living gastropods species is estimated to exceed 100,000 (Appeltans et al. 2012), accounting for approximately 80% of all extant molluscs. Gastropods exhibit very broad distribution ranges encompassing a wide range of ecological settings, ranging from the Himalayas at altitudes of 5,500 meters to oceanic trenches exceeding 10,000 meters in depth (Annandale and Rao 1925; Glöer and Bössneck 2013). Fossil records indicate that gastropods originated from shallow water environments during the early Cambrian (Parkhaev 2017). Currently, approximately 45–50% of gastropod species are marine, while 40–45% are terrestrial, and about 5% live in freshwater habitats (Krug et al. 2022), making it a great model for studying the molecular evolution linked with habitat transitions.

Frequent transitions across marine, freshwater, and terrestrial habitats as seen in gastropods is exceptional among animals. Previous studies have demonstrated that gastropods colonized freshwater habitats 33-38 times independently, including lineages of Caenogastropoda, Heterobranchia, and Neritimorpha (Strong et al. 2008). Furthermore, transitions to the freshwater have frequently served as a vital evolutionary ‘stepping stone’, facilitating 14 of the 30 known independent transitions to terrestrial habitats (Vermeij and Watson-Zink 2022). Similarly, the colonization of ‘extreme’ deep-sea environments such as hydrothermal vents by extant gastropods dates back to the Mesozoic, as multiple lineages including all six subclasses, most notably Neomphaliones with a large number of endemic lineages, have independently colonized chemosynthetic ecosystems (Vrijenhoek 2013; Zhang et al. 2024).

Chromosomal rearrangements are increasingly recognized as powerful engines of evolutionary change that can facilitate adaptation by reshaping gene linkages and regulatory landscapes, including during habitat transitions. A recent study on annelids demonstrated that an episodic burst of chromosomal rearrangements was intimately linked to the origin of non-marine lineages, suggesting that the reshaping of the genomic landscape play a pivotal role in overcoming environmental barriers (Vargas-Chávez et al. 2025). Across diverse taxa, structural variations act as a ubiquitous adaptive mechanism. For instance, chromosomal inversions drive local ecotype adaptation in plants (Lowry and Willis 2010), preserve adaptive divergence during ecological speciation in insects (Fuller et al. 2019), and help maintain distinct ecological niches in vertebrates (Kirubakaran et al. 2016). These examples underscore chromosomal rearrangements as a fundamental mechanism for environmental adaptation across the tree of life. However, despite our extensive knowledge of the numerous ecological transitions seen in gastropods, the link between these transitions and genomic rearrangements have not been studied.

Reconstructing ancestral karyotype compositions of Gastropoda and each of its subclasses are fundamental to understanding the extent of subsequent structural modifications that coincide with habitat shifts. However, there have been significant incongruencies in the relationships among the six subclasses derived from fossil evidence and morphology, as well as those from various molecular phylogenetic reconstructions using a range of different data types (Bieler 1992; Ponder and Lindberg 2008; Kocot et al. 2011; Zapata et al. 2014; Cunha and Giribet 2019; Uribe et al. 2022; Zhong et al. 2022; Chen et al. 2025b). Most notably, morphology and fossil evidence suggest Patellogastropoda is the sister of all other subclasses (i.e. the Orthogastropoda hypothesis) (Ponder and Lindberg 2008), but most molecular data point to Patellogastropoda being sister to Vetigastropoda + Neomphaliones (the Psilogastropoda hypothesis) (Zapata et al. 2014; Sun et al. 2020; Chen et al. 2025b). Such conflicting topologies have direct implications for inferring the evolutionary history of the ancestral gastropod karyotype. Recently, chromosome-level macrosynteny has emerged as a potential additional phylogenetic tool for resolving deep nodes (Schultz et al. 2023; Schultz et al. 2026), since large-scale chromosomal rearrangements are rare events with exceptionally low homoplasy. Therefore, exploring macrosyntenic patterns could provide another line of evidence to resolve the gastropod phylogeny and then accurately reconstruct their ancestral karyotype.

A key limiting factor to resolving the chromosomal evolution across different gastropod subclasses is that most of the available chromosome-level genomes are from Caenogastropoda and Heterobranchia (Chen et al. 2025b). Here, we present high-quality chromosome-level genomes of five gastropod species from the other four subclasses, and use these new data in combination with existing genomes to reconstruct the ancestral karyotypes. These included the freshwater *Vittina coromandeliana* and the deep-sea vent-endemic *Shinkailepas gigas* from Neritimorpha, the shallow-water turbinid *Turbo cornutus* from Vetigastropoda, the vent-endemic neomphalid *Neomphalus fretterae* from Neomphaliones, and *Cellana toreuma* from Patellogastropoda. Through macrosynteny analyses, we aimed to elucidate the evolutionary trajectories from the ancestral gastropod karyotype to the extant karyotypes of each subclass, and to evaluate the functional significance of chromosomal rearrangements in the adaptation of freshwater and deep-sea species.

## Materials and Methods

### Sample collection

The neritid *Vittina coromandeliana* (Neritimorpha) is a common species in the aquarium trade, and specimens were purchased from a local aquarium market in Qingdao, China and morphologically identified. The phenacolepadid *Shinkailepas gigas* (Neritimorpha) was collected using a suction sampler on the human-occupied vehicle (HOV) *Shinkai 6500* during dive #1770 on-board R/V *Yokosuka* from the Mokuyo Seamount hydrothermal vent field, Izu-Ogasawara Arc (28°18.9088’N, 140°33.8646’E) at a depth of 1254 m (Chen et al. 2025a). Specimens of the turbinid snail *Turbo cornutus* (Vetigastropoda) were collected on a rocky shore during the low tide at Zhoushan Island, Zhejiang Province, China. The neomphalid *Neomphalus fretterae* (Neomphaliones) was sampled during dive #606 of the remotely operated vehicle (ROV) *SuBastian* on-board R/V *Falkor (too)* at the Zombie vent (0°46.2720′N, 85°55.9920′W), Tempus Fugit vent field, Galápagos Rift, at a depth of 2514 m (Chen et al. 2024). The nacellid *Cellana toreuma* (Patellogastropoda) specimens were collected from a rocky shore at Taipingjiao, Qingdao, China during low tide.

### Genome sequencing

Genomic DNA from foot tissue of each species was extracted using the Magnetic Universal Genomic DNA Kit (Tiangen Biotech (Beijing) Co., Ltd.). For *V. coromandeliana*, *T. cornutus*, and *C. toreuma*, short-insert (350 bp) paired-end libraries were constructed with NEBNext® Ultra™ DNA Library Prep Kit for Illumina (NEB) and sequenced on an Illumina Hiseq XTen platform with the read length of 150 bp. For *V. coromandeliana*, *T. cornutus*, *C. toreuma*, and *S. gigas*, the Oxford Nanopore Technology (ONT) libraries were constructed using Ligation Sequencing Kit (SQK-LSK109) and then sequenced on a PromethION DNA sequencer (Oxford Nanopore Technologies, UK). For *S. gigas* and *N. fretterae*, SMRTbell libraries were constructed using SMRTbell Express Template Prep Kit 2.0 (Pacific Biosciences, Menlo Park, CA, USA) and the libraries were sequenced on a PacBio Sequel II platform (Pacific Biosciences) in the CCS (circular consensus sequencing) mode.

### Hi-C Sequencing

In order to obtain chromosome-level genome assemblies, genomes of the five species were further sequenced on a Hi-C platform. Hi-C libraries were constructed with the standard protocol (Belton et al. 2012) with certain modifications. Samples were ground with liquid nitrogen and cross-linked with 4% formaldehyde, followed by chromatin digestion using the restriction endonuclease MboI. The resulting libraries were then sequenced on an Illumina Hiseq platform with 150 bp paired-end reads.

### Transcriptome sequencing

For transcriptome sequencing, digestive gland, foot muscle, and mantle were dissected from *V. coromandeliana* and *C. toreuma*. Digestive gland, foot muscle, mantle, gill, and the operculum-secreting tissue were dissected from *T. cornutus*. Digestive gland, foot muscle, mantle, gill, and the head were dissected from *S. gigas*. Digestive gland, foot muscle, mantle, gill, and gonad were dissected from *N. fretterae*. RNA sequencing libraries were prepared using the NEBNext® Ultra™ RNA Library Prep Kit for Illumina® (NEB) and sequenced on an Illumina HiSeq platform.

### Genome assembly

Raw ONT data in the *.fast5* format were base called using Guppy v.5.0.11 with the configure file of dna_r9.4.1_450bps_sup_prom.cfg and SUP mode. For *V. coromandeliana*, *T. cornutus* and *C. toreuma*, the genomes were assembled by Shasta v.0.7.0 (Shafin et al. 2020), Shasta v.0.8.0, Raven v.1.5.1 (Vaser and Šikić 2020), Flye v.2.9-b1768 (Kolmogorov et al. 2020), and NextDenovo v.2.4.0 (Hu et al. 2024b) using ONT reads which were sub-sampled with different cutoff lengths using Seqtk v.1.3-r106. Following comparison of assembly statistics of different pipelines, the genomes assembled from the NextDenovo pipeline for *C. toreuma* and *V. coromandeliana*, and the Shasta pipeline for *T. cornutus* were the best and thus used for downstream analyses. The assembled contigs were error-corrected twice using the ONT fastq data with length over 3Kb by Flye v.2.9-b1768, and heterozygous contigs were removed using Purge Dups v.1.2.5 (Guan et al. 2020). Then, the resultant contigs were polished twice with MaSuRCA v.4.0.5 (Zimin et al. 2013) using Illumina reads. For *S. gigas*, the genome was assembled by NextDenovo v.2.4.0 with ONT reads. Haplotypic duplication was removed using Purge-dups v.1.2.5, and the resultant contigs were polished with NextPolish2 v.0.2.1 (Hu et al. 2024a) using PacBio HiFi reads. For *N. fretterae*, the genome was assembled using hifiasm v.0.25.0-r726 (Cheng et al. 2021) with PacBio HiFi reads, followed by the removal of heterozygous contigs using Purge_dups v.1.2.5.

Genome assembly completeness was monitored at each step using Benchmarking Universal Single Copy Orthologs (BUSCO) v.5.2.2 (Simão et al. 2015) with the odb10 metazoan dataset. Bacterial sequence contaminations in the assembled contigs were checked by BLASTP searching against the NCBI NT database, and then confirmed by BlobTools v.1.1.1 (Laetsch and Blaxter 2017) and MEGAN 6. Detected bacterial sequences were removed from the assembled genomes.

### Genome scaffolding

Adapters and low-quality Hi-C raw Illumina reads were removed by Trimmomatic v.0.39 (Bolger et al. 2014). The Hi-C reads were mapped to the assembled contigs and filtered with HiC-Pro v.3.1.0 (Servant et al. 2015). Based on the filtered, valid Hi-C reads, the Juicer v.1.6 (Durand et al. 2016) pipeline was used to further process the interaction data and generate Hi-C contact matrices. Then, genomic scaffolding was conducted by 3D-DNA v.180922 (Dudchenko et al. 2017). Several manual corrections were performed in Juicebox.

### Gene model prediction and annotation

Adapters of raw RNA Illumina reads were removed by Trimmomatic v.0.39. Then transcriptomes were assembled by two versions: one de novo assembled by Trinity v.2.13.2 (Haas et al. 2013); another assembled by Trinity v.2.13.2 under the genome-guided mode using *.bam* file generated by aligning to the genome with histat2 v.2.2.1 (Kim et al. 2019). These two versions of assembled transcriptome were merged and further clustered using cd-hit-est v.4.8.1 (Li and Godzik 2006) with a minimum sequence identity of 0.97. Protein sequences of molluscs from the SWISS-PROT database and NCBI were downloaded and used to help guide the prediction process.

Repetitive sequences of the genomes were identified using a two-step strategy. First, de novo species-specific repeat libraries were built by RepeatModeler v.2.0.2a, and RepeatMasker v.4.1.2 was further run against the de novo species-specific library for each genome. Subsequently, RepeatMasker v.4.1.2 was used to search repbase (RepBase v.20181026) to identify homologous repeats. Then, contigs smaller than 100 Kb in the repeats-masked genomes were removed. Raw RNA reads of one individual were trimmed by Trimmomatic v.0.39 and then the reads were aligned to the genome using histat2 v.2.2.1 which generated *.bam* file. Then, the repeats-masked genomes and the *.bam* file were used to train Augustus v.3.4.0 (Stanke et al. 2006) using BRAKER v.2.1.6 (Brůna et al. 2021) for de novo gene prediction.

Gene model prediction was performed using Maker v.3.01.04 based on their evidence-based weights with three combined evidences, including transcripts-based evidence, protein-evidence and Augustus gene predictions. Gene models were also predicted by EVidence Modeler v.1.1.1 (Haas et al. 2008), integrating evidence from transcriptome alignments, protein alignments, and ab initio predictions. For transcriptome alignments, transcriptomes were mapped to the five genomes in the PASA pipeline using the BLAT v.35.1 mapping tools. For protein alignments, protein sequences of molluscs from the SWISS-PROT database and NCBI were downloaded and aligned to genomes. Ab initio predictions were obtained from training Augustus v.3.4.0 using BRAKER v.2.1.6 for gene de novo prediction. Then, the structural annotations of genes were updated to add UTRs using the annotation update method in the PASA pipeline.

### Phylogenomic reconstruction

Phylogenomic reconstruction was constructed using the VEHoP v.1.3 pipeline (Li et al. 2026) with occupancy threshold of 82%. Maximum likelihood (ML) reconstruction was conducted with IQ-TREE v.2.4.0 (Nguyen et al. 2014). For ML, the optimal models for each partition were selected with the setting of “-m MFP”. A posterior mean site frequency (PMSF) model was also performed with the setting of “-m LG+C60+F+Γ”, using the guide tree generated from MFP model. The topological support was evaluated using 1,000 ultrafast bootstrap replicates.

Divergence time was estimated utilizing the MCMCTREE program implemented in PAML v4.8 (Pagel and Meade 2006; Reis and Yang 2011). The time-calibration incorporated three constraints derived from fossil records and geological events: (1) the first appearance of molluscs between a hard minimum of 532 Ma and a hard maximum of 549 Ma (Benton et al. 2015); (2) the divergence between Caenogastropoda and Heterobranchia with a hard minimum of 390 Ma (Jörger et al. 2010); and (3) a hard minimum age of 132.9 Ma for the origin of Neomphalidae based on the earliest reliable fossil record dating back to the Early Cretaceous (Valanginian) (Kiel and Campbell 2005; Campbell et al. 2008).

### Identification of ancestral linkage groups

The identification of ancestral linkage groups (ALGs) was performed following previously established methodologies (Simakov et al. 2020; Simakov et al. 2022; Schultz et al. 2023; Sigwart et al. 2024). To infer the ancestral and conserved ALGs for Gastropoda, six chromosome-level genome assemblies were utilized, each representing one of the six gastropod subclasses: *Alviniconcha adamantis* (Caenogastropoda), *Berghia stephanieae* (Heterobranchia), *Vittina coromandeliana* (Neomphaliones), *Turbo cornutus* (Vetigastropoda), *Chrysomallon squamiferum* (Neomphaliones), and *Cellana toreuma* (Patellogastropoda). Furthermore, ALGs for each gastropod subclasses were inferred using the following genomes: *Alviniconcha adamantis*, *Littorina brevicula*, and *Pomacea canaliculata* for Caenogastropoda; *Biomphalaria glabrata*, *Berghia stephanieae*, and *Elysia timida* for Heterobranchia; *Vittina coromandeliana* and *Shinkailepas gigas* for Neritimorpha; *Turbo cornutus*, *Gibbula magus*, and *Haliotis asinina* for Vetigastropoda; *Chrysomallon squamiferum*, *Gigantopelta aegis*, and *Neomphalus fretterae* for Neomphaliones; and *Cellana toreum*a and *Patella vulgata* for Patellogastropoda.

First, all-against-all protein sequence alignments were executed between genome pairs via DIAMOND blastp (Buchfink et al. 2021). Genes were classified as putative orthologs if they exhibited a mutual best hit (MBH) relationship within each species pair. Subsequently, linkage relationships of orthologous genes were established across multiple genomes. Only strict single-copy orthologs, which are defined as homologous genes present in all genomes within a specific comparison group, were retained. Finally, the statistical significance of linkage groups between any two genomes was assessed using Fisher’s exact test implemented in the R package macrosyntR (El Hilali and Copley 2023), with a significance threshold set at adjusted *P* < 0.001.

### Macrosynteny analysis

Macrosynteny analysis was conducted to investigate chromosomal rearrangements within each subclass. Pairwise protein alignments were first performed using DIAMOND blastp. Gene pairs exhibiting mutual best-hit (MBH) relationships were identified as putative orthologs. To establish a reference framework for comparative analyses, a subset of genes was subsequently extracted for each species, consisting of MBH genes assigned to the ancestral linkage groups of the corresponding subclass. Finally, macrosyntenic relationships were visualized by constructing comparative chromosomal diagrams using the R package macrosyntR.

### Identification of topologically associating domains and characterization of chromosomal rearrangements

To investigate the potential association between evolutionary chromosomal rearrangement events and the three-dimensional (3D) genomic architecture, topologically associating domains (TADs) were systematically identified based on Hi-C data to determine whether structurally rearranged regions were constrained within TAD internal boundaries (Zhou et al. 2026). First, the HiC-Pro pipeline was used to process species-specific Hi-C sequencing data, constructing an index via in silico restriction enzyme digestion, mapping filtered high-quality reads to the reference genome, and removing invalid interactions to generate genome-wide matrices. Raw interaction matrices were extracted at a 40 kb resolution for high-precision 3D structural resolution, followed by the extraction of local chromosomal interactions using a custom script and conversion into a multi-scale .cool format using toCooler. The hitad algorithm from the TADLib package was then applied to precisely identify TADs and their boundaries at 40 kb resolution, while insulation scores were calculated using the fanc software across multiple sliding windows (1.0 Mb to 4.0 Mb) to quantitatively define highly conserved TAD boundaries as regions with lower insulation scores (stronger insulation). Finally, standardized matrices were converted to .h5 format via HiCExplorer (Wolff et al. 2022), and high-resolution genomic tracks (Hi-C heatmaps, insulation scores, and TAD structures) were plotted using pyGenomeTracks; the coordinates of regions exhibiting syntenic breaks and chromosomal rearrangements were then cross-mapped with physical TAD boundary coordinates to systematically assess whether evolutionary genomic rearrangements occurred conservatively within a single TAD or disrupted original boundary structures.

### KEGG pathway coverage analysis of rearranged genes

Functional annotation of the rearranged genes (i.e., genes located within genomic regions that underwent structural changes, such as fissions or fusions, relative to the ancestral karyotype) was performed using the BlastKOALA web server to assign KEGG (Kyoto Encyclopedia of Genes and Genomes) Orthology (KO) identifiers based on sequence similarity. Subsequently, a custom R script using the dplyr package was employed to conduct pathway coverage analysis. The coverage for each pathway was calculated as the ratio of the number of rearranged genes mapped to that specific pathway to the total number of background genes annotated to the same pathway, expressed as a percentage.

To ensure biological relevance and statistical reliability, the raw results were subjected to filtering criteria. Pathways annotated as “General function” were excluded from the dataset, and only pathways containing at least two rearranged genes were retained for further analysis. The top 20 pathways were selected based on coverage ranking and gene count. Finally, visualization of the results was conducted using the R package ggplot2, where the pathway coverage and gene counts were represented in a bubble plot.

## Results

### Chromosome-level assembly and annotation of the five gastropod genomes

Chromosome-level genome assemblies and comprehensive gene annotations for the five gastropod species were successfully generated (Fig. 1). Specifically, the genome sizes of *V. coromandeliana*, *S. gigas*, *T. cornutus*, *N. fretterae*, and *C. toreuma* were 1.51 Gb, 1.07 Gb, 1.77 Gb, 958.21 Mb, and 325.96 Mb, with contigs successfully anchored to 14, 14, 18, 15, and 9 chromosomes, respectively (Fig. S1). Based on transcript-based evidence, protein homology, and *ab initio* predictions, 26,967, 26,144, 27,575, 26,496, and 20,804 gene models were identified for *V. coromandeliana*, *S. gigas*, *T. cornutus*, *N. fretterae*, and *C. toreuma*, respectively. The completeness of these annotated gene models was further validated by BUSCO scores of 93.4%, 89.9%, 93.5%, 96.4%, and 95.6%, respectively.

**Figure 1.**
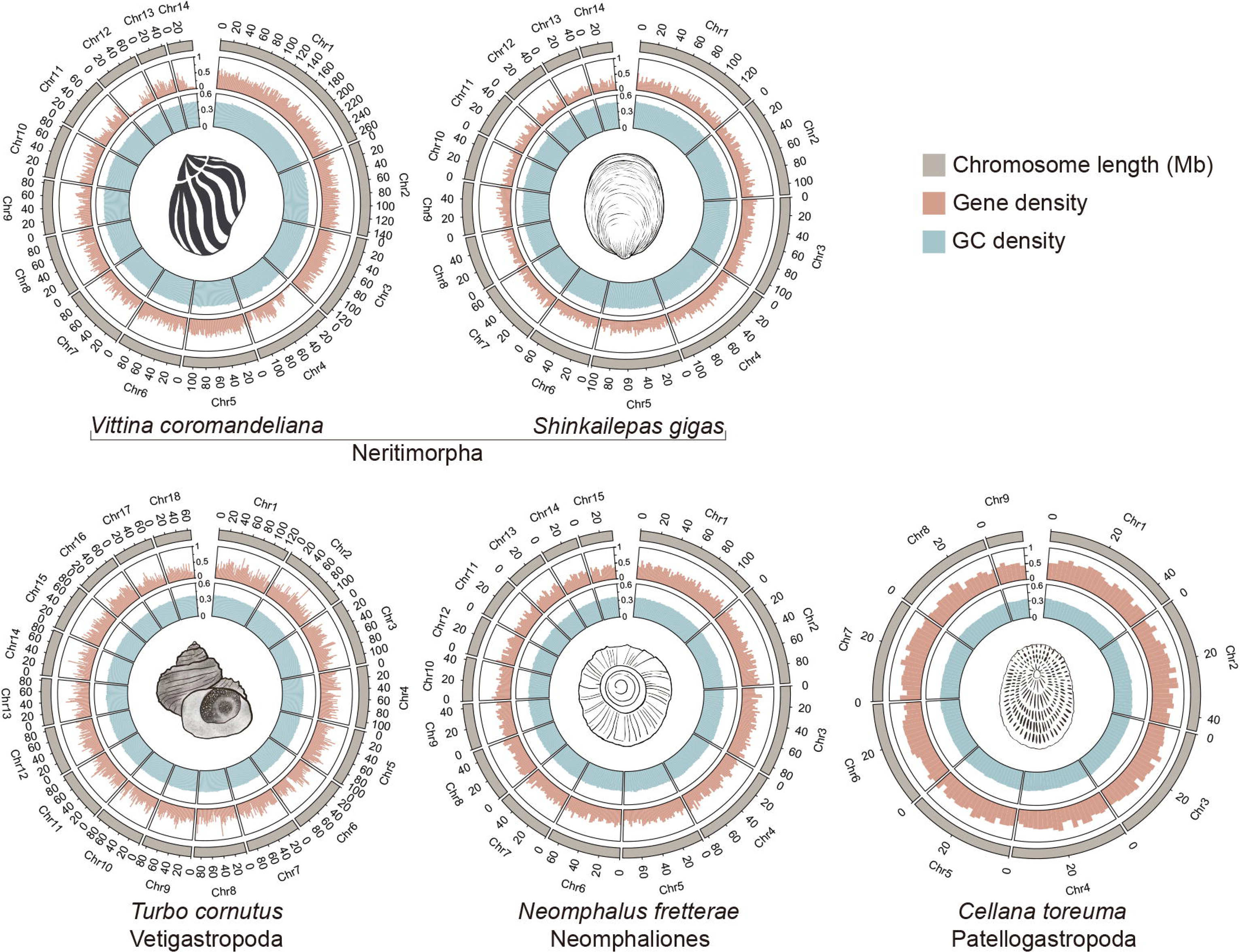
Circos plots characterizing the genomic landscapes of the five gastropod species with their chromosome-level genomes sequenced anew: *Vittina coromandeliana* (Neritimorpha), *Shinkailepas gigas* (Neritimorpha), *Turbo cornutus* (Vetigastropoda), *Neomphalus fretterae* (Neomphaliones), and *Cellana toreuma* (Patellogastropoda). The tracks, arranged from the outer to the inner circles, represent: Chromosome length (Mb); Gene density; and GC content density.

### Ancestral linkage groups for each gastropod subclass

The ancestral gastropod karyotype is reconstructed to comprise 20 ancestral gastropod linkage groups (GLGs). These 20 GLGs exhibit strict one-to-one relationship with the ancestral molluscan linkage groups (MLGs) (Fig. S2). Comparative analysis across the six gastropod subclasses revealed subsequent numerical variation in linkage groups driven by chromosomal rearrangements (Fig. 3). The ancestral Caenogastropoda linkage groups (CgLGs), Heterobranchia linkage groups (HbLGs), and Vetigastropoda linkage groups (VgLGs) each retained 18 linkage groups. This reduction from GLGs is characterized by specific events where ancestral linkage groups undergo fusion-with-mixing to form a single chromosome. For instance, GLG 1 and GLG 18 underwent fusion-with-mixing to form one chromosome, while GLG 19 and GLG 20 similarly went through fusion-with-mixing to form another chromosome. Despite these chromosomal rearrangements, these 18 linkage groups maintained significant syntenic conservation.

The remaining three subclasses exhibited a more pronounced reduction in linkage groups involving further fusion-with-mixing events. The ancestral Neomphaliones linkage groups (NpLGs) comprised 15 linkage groups, a state achieved through specific chromosomal rearrangements such as GLG 12, 13, and 10 undergoing fusion-with-mixing to form a chromosome. The ancestral Neritimorpha linkage groups (NmLGs) contained 14 linkage groups resulting from extensive chromosomal rearrangements, notably GLG 8, 11, 9, 13, and 14 underwent fusion-with-mixing to form consolidated chromosomes. The ancestral Patellogastropoda linkage groups (PgLGs) displayed the most reduced chromosomal numbers among the described groups, with a total of 10 linkage groups. This configuration emerged from widespread fusion-with-mixing that effectively halved the ancestral count by consolidating the 20 GLGs into 10 PgLGs. Figure 3 visualizes these syntenic relationships and the evolutionary transitions from the 20 GLGs to the divergent chromosomal architecture characteristic of the six subclasses.

### Macrosynteny evidence for Psilogastropoda vs Orthogastropoda hypotheses

Chromosomal rearrangement scenarios were tested on two distinct topologies: Tree 1 based on the whole-genome tree of Chen et al. (2025) supporting Psilogastropoda, and Tree 2 based on the mitogenome tree of Uribe et al. (2019) which supports Orthogastropoda (Fig. 2). Under the Tree 1 topology for Psilogastropoda, the evolution of the Patellogastropoda karyotype (PgLGs) from ancestral gastropod linkage groups (GLGs) followed a two-step pathway. The first step involved the transition from GLGs to the common ancestor of VgLGs, NpLGs, and PgLGs (Vg_Np_Pg LGs), which maintained a stable one-to-one relationship, while the second step from the Vg_Np_Pg LGs to PgLGs was characterized by significant rearrangements. Specifically, apart from PgLG10 which showed a one-to-one relationship with Vg_Np_Pg LG17, the remaining nine PgLGs were formed through extensive fusion-with-mixing events. Conversely, in the Tree 2 topology for Orthogastropoda, the PgLGs evolved directly from the GLGs; similar to the second step of Tree 1, this process was defined by a one-to-one relationship between PgLG10 and GLG17, while the other nine PgLGs resulted from large-scale fusion-with-mixing.

**Figure 2.**
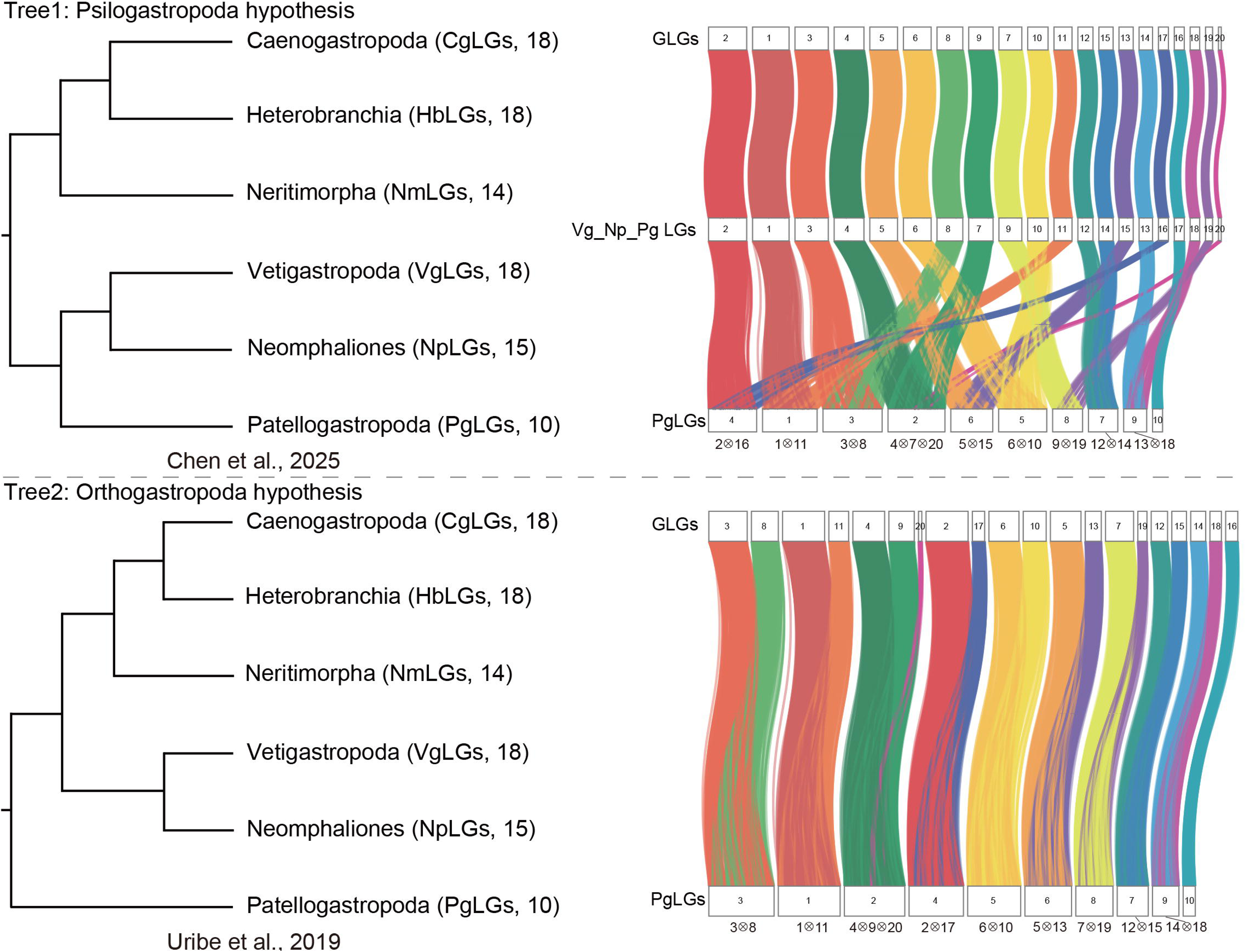
Comparison of chromosomal evolutionary trajectories from ancestral Gastropod Linkage Groups (GLGs) to Patellogastropoda Linkage Groups (PgLGs) under two contrasting topologies, with the chromosomal evolutionary paths reconstructed based on the topologies. Tree 1 (Psilogastropoda hypothesis) suggests a two-stage process: GLGs first evolves into the common ancestor of Vetigastropoda, Neomphaliones, and Patellogastropoda (named as Vg_Np_Pg LGs), followed by the evolution into PgLGs. Tree 2 (Orthogastropoda hypothesis) proposes a single-stage direct evolution from GLGs to PgLGs. The diagrams on the right illustrate the chromosomal synteny.

**Figure 3.**
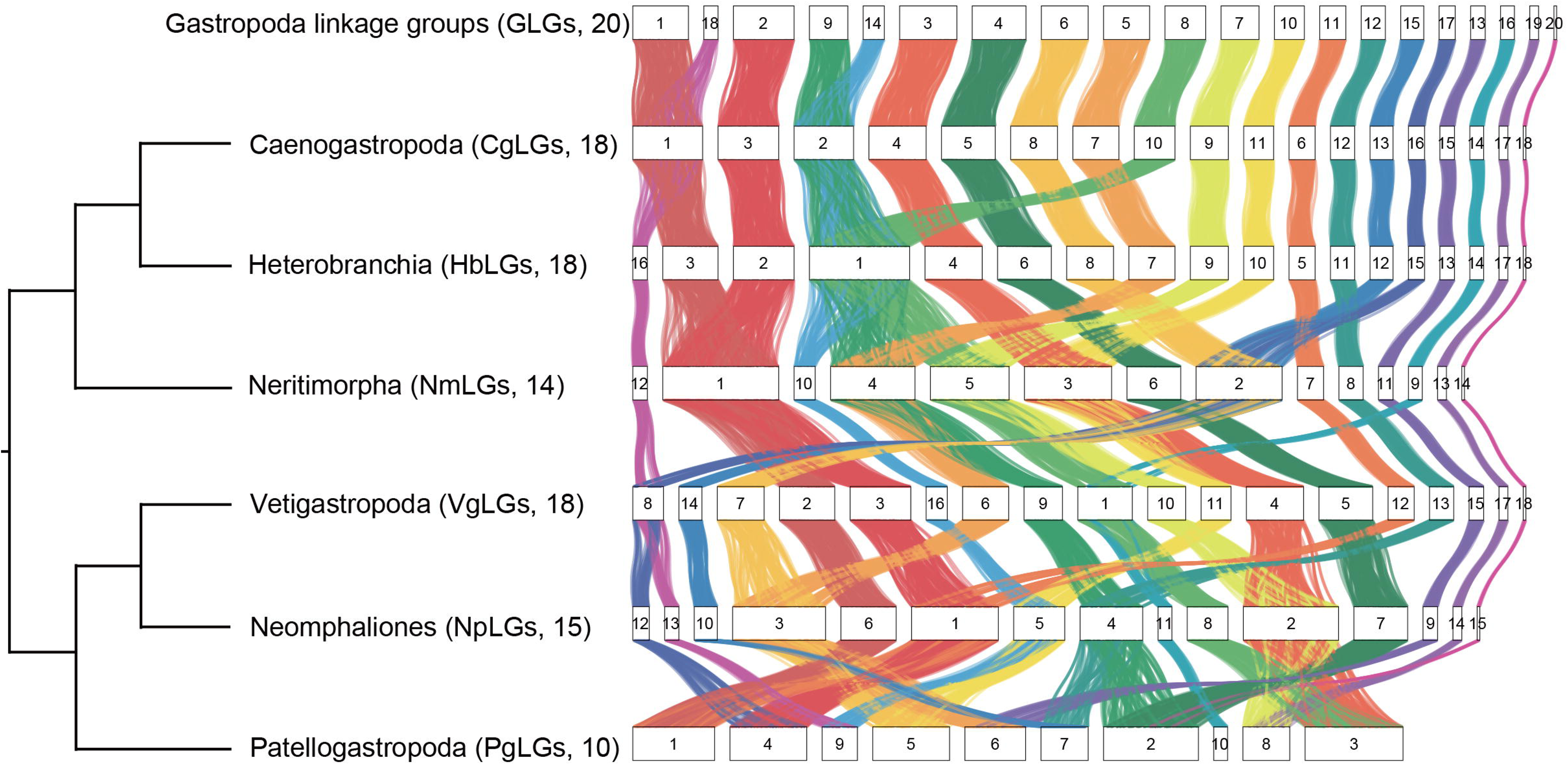
Reconstruction and synteny evolution of ancestral linkage groups (ALGs) across the six gastropod subclasses. The diagram illustrates the chromosomal evolutionary relationships starting from the ancestral Gastropoda Linkage Groups (GLGs, n=20). The reconstructed ancestral karyotypes for the six subclasses are shown as follows: Caenogastropoda (CgLGs, n=18), Heterobranchia (HbLGs, n=18), Neritimorpha (NmLGs, n=14), Vetigastropoda (VgLGs, n=18), Neomphaliones (NpLGs, n=15), and Patellogastropoda (PgLGs, n=10).

### Chromosomal rearrangements linked with habitat transitions within gastropod subclasses

In the subclass Caenogastropoda, chromosomal rearrangements showed a strong correlation with habitat transitions compared to CgLGs (Fig. 4). Freshwater species exhibited the most significant chromosomal rearrangements, with *Pomacea canaliculata* going through three chromosomal fusions, while *Semisulcospira habei* displayed extensive end-to-end fusions involving seven out of eight chromosomes. Among the shallow-water marine species, *Littorina brevicula* showed only a single end-to-end fusion, whereas members of the order Neogastropoda such as *Monoplex corrugatus*, *Babylonia areolata*, *Conus ventricosus*, *Rapana venosa*, and *Stramonita haemastoma* have experienced a partial genome duplication in their common ancestor (Farhat et al. 2023). In contrast, the karyotype of the deep-sea vent-endemic species *Alviniconcha adamantis* remained highly conservative, maintaining a one-to-one relationship with CgLGs.

**Figure 4.**
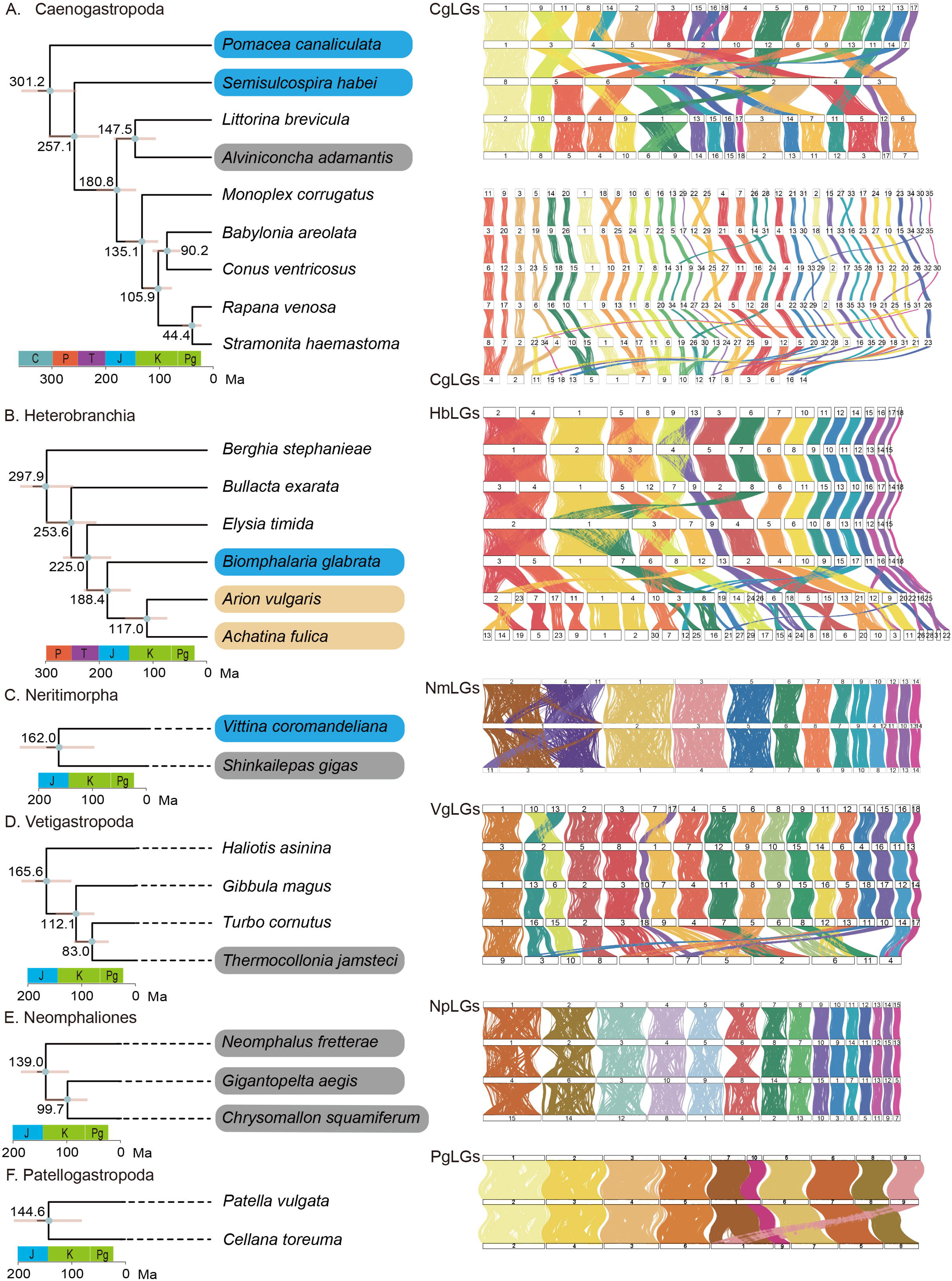
Phylogenetic relationship and chromosomal synteny patterns across five gastropod subclasses, showing evolutionary relationships and chromosomal rearrangements within the subclasses (A) Caenogastropoda, (B) Heterobranchia, (C) Neritimorpha, (D) Vetigastropoda, and (E) Neomphaliones. The left panels display time-calibrated phylogenetic trees with divergence times (Ma) indicated at the nodes. The right panels show chromosomal synteny between the reconstructed ancestral linkage groups (CgLGs, HbLGs, NmLGs, VgLGs, and NpLGs) and the genomes of the extant species. Species are color-coded based on their habitats to highlight ecological adaptations: shallow water (white without highlight), freshwater (blue), terrestrial (orange), and deep sea (grey).

Within Heterobranchia, the evolutionary stability of chromosomes varied across environmental niches relative to HbLGs (Fig. 4). Terrestrial species, specifically *Arion vulgaris* and *Achatina fulica*, are characterized by whole-genome duplications. In contrast, the freshwater species *Biomphalaria glabrata* maintained a conservative one-to-one relationship with HbLGs. Shallow-water marine species exhibited diverse patterns, where the karyotype of *Bullacta exarata* remained conservative but both *Berghia stephanieae* and *Elysia timida* have undergone small-scale fusion-with-mixing events involving three chromosomes each.

The subclasses Neritimorpha and Vetigastropoda also displayed habitat-specific chromosomal dynamics (Fig. 4). In Neritimorpha, the deep-sea species *Shinkailepas gigas* showed a strict one-to-one relationship with NmLGs. However, the freshwater species *Vittina coromandeliana* exhibited one mixed fusion and four chromosomes derived from ancestral fissions. Similarly, in Vetigastropoda, shallow-water species like *Gibbula magus* and *Turbo cornutus* had conserved karyotypes relative to VgLGs, although *Haliotis asinina* showed two end-to-end fusions. Notably, the deep-sea vent-endemic vetigastropod *Thermocollonia jamsteci* underwent extensive rearrangements, with seven of its eleven chromosomes resulting from fusion-with-mixing. Finally, all examined species in Neomphaliones, including *Neomphalus fretterae*, *Gigantopelta aegis*, and *Chrysomallon squamiferum*, are all deep-sea vent species that exhibited remarkable stability, all maintaining a one-to-one relationship with the ancestral NpLGs.

### Association analysis between chromosomal rearrangements and 3D genomic TAD structures

To explore how evolutionary genomic structural variations in gastropods are constrained by their 3D spatial architecture, the physical positions of the identified chromosomal rearranged regions were cross-referenced with TADs. Quantitative statistical analysis of the positions of rearranged genes in three representative species, *Semisulcospira habei*, *Vittina coromandeliana*, and *Thermocollonia jamsteci*. These species were selected because their relatively lower degrees of chromosomal fusion-with-mixing enable robust statistical evaluation, avoiding the highly complex fusion-with-mixing observed in other gastropods that confound spatial comparisons. The analysis revealed a significant spatial preference for genomic rearrangement events to distribute within TADs. Across the analyzed species, the vast majority of rearranged genes are located in the interior of TADs rather than at the boundaries. Specifically, up to 95.8% (23/24) of the rearranged genes in *S. habei* are located within the TAD interior (with only 1 gene at the boundary), while the proportion of internally rearranged genes reached 89.5% (51/57) in *T. jamsteci* and 78.0% (39/50) in *V. coromandeliana*. A graphical representation of these spatial preferences is available on Figshare (DOI: 10.6084/m9.figshare.33403453).

### Pathway coverage analysis of rearranged genomic regions

To identify the functional drivers of chromosomal evolution across habitat transitions, KEGG pathway coverage analyses were performed on genes within rearranged genomic regions (Fig. 5). In freshwater species, *Semisulcospira habei* (Caenogastropoda) displays extensive end-to-end fusions, while *Biomphalaria glabrata* (Heterobranchia) maintains a conservative one-to-one relationship with HbLGs. Consequently, *Pomacea canaliculata* (Caenogastropoda) and *Vittina coromandeliana* (Neritimorpha) were selected for analysis to compare their adaptive mechanisms.

**Figure 5.**
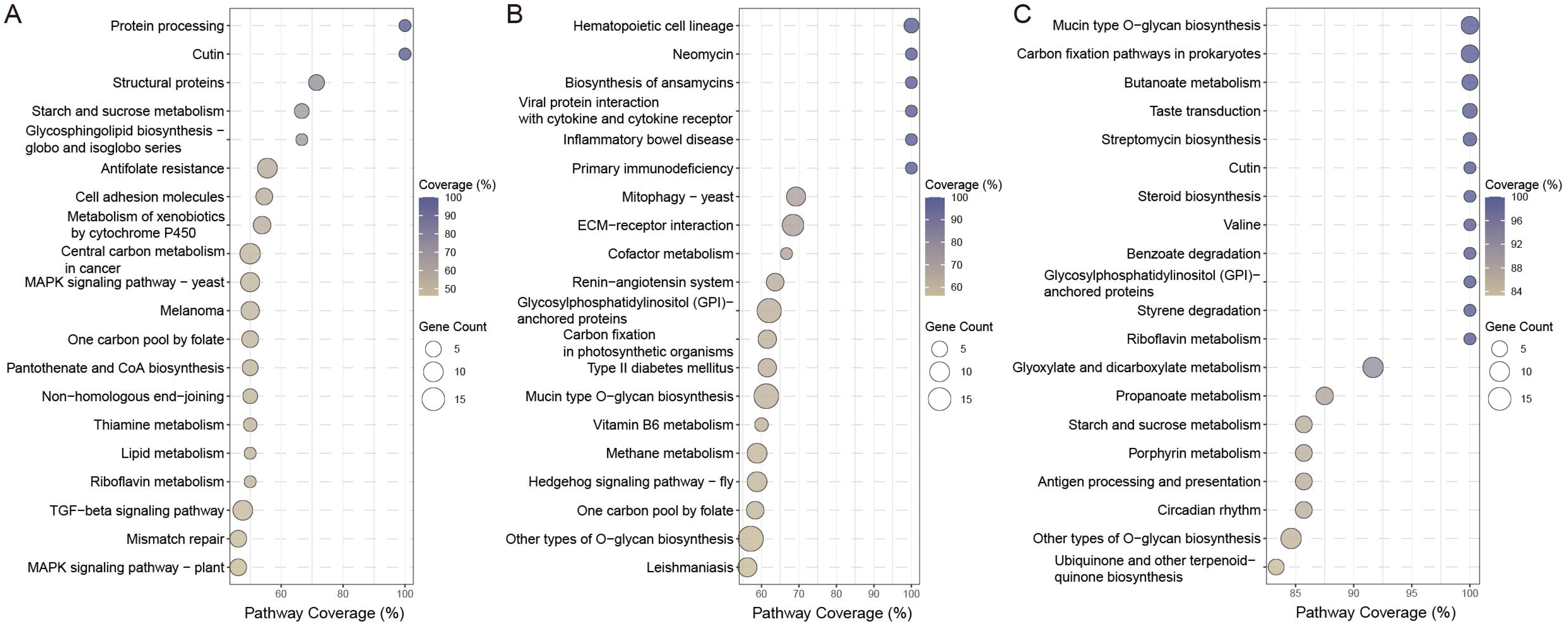
KEGG pathway coverage analysis of genes associated with chromosomal rearrangements. The dot plots illustrate the functional enrichment of genes located in rearranged chromosomal regions across different lineages. (A) Analysis of genes involved in rearrangements in the freshwater caenogastropod species *Pomacea canaliculata*. (B) Analysis of genes involved in rearrangements in the freshwater neritimorph species *Vittina coromandeliana*. (C) Analysis of genes associated with chromosomal rearrangements identified between the deep-sea endemic Neomphaliones Linkage Groups (NpLGs) and ancestral Gastropod Linkage Groups (GLGs). The x-axis represents the percentage of pathway coverage. The size of each dot corresponds to the number of genes annotated to the pathway (Gene Count), and the color gradient indicates the coverage percentage.

In *P. canaliculata*, the rearranged regions were significantly associated with starch and sucrose metabolism, reflecting an efficient conversion of aquatic plant nutrients to support rapid growth and spawning. Additionally, high coverage in the metabolism of xenobiotics by cytochrome P450 and the MAPK signaling pathway highlighted enhanced detoxification capabilities and sensory resilience to extreme physical stressors. Conversely, *V. coromandeliana* prioritized physiological homeostasis through its rearranged regions, featuring pathways such as hematopoietic cell lineage and viral protein interaction with cytokine receptors for advanced biological defense, as well as the renin-angiotensin system for osmotic regulation. The enrichment of mucin-type O-glycan biosynthesis and ECM-receptor interaction further suggests a reinforced protective barrier against chemical irritation and physical river scouring (Davies and Hawkins 1998; Liegertová and Malý 2023).

Among deep-sea hydrothermal vent species, while *Alviniconcha adamantis* (Caenogastropoda) and *Shinkailepas gigas* (Neritimorpha) maintained highly conservative one-to-one relationships with their respective ancestral linkage groups, *Thermocollonia jamsteci* (Vetigastropoda) has undergone extensive chromosomal rearrangements. The NpLGs relative to the ancestral GLGs were used to capture the broad functional shifts associated with deep-sea colonization. Several ecologically specialized pathways showed remarkable coverage in these rearranged regions. The mucin-type O-glycan biosynthesis pathway reached 100% coverage, likely facilitating a high-strength mucus barrier against the acidic and corrosive hydrothermal vent fluids. Furthermore, steroid biosynthesis and multiple detoxification pathways suggested mechanisms to stabilize cell membranes against extreme hydrostatic pressure while neutralizing heavy metal toxins. Collectively, chromosomal rearrangements in deep-sea hydrothermal lineages were linked to chemosymbiosis with multi-layered physiological safeguards, enabling homeostatic control in extreme environments.

## Discussion

The reconstruction of the ancestral Gastropod Linkage Groups (GLGs) offered a novel chromosomal perspective on the early radiation of this class, but also highlighted the limitations of macrosynteny in resolving deep nodes of the phylogeny. We attempted to test the support for the Psilogastropoda vs Orthogastropoda hypotheses of the gastropod tree with macrosynteny by evaluating the path required from ancestral GLGs to Patellogastropoda Linkage Groups (PgLGs), to see which is more parsimonious. However, the macrosynteny analysis revealed that both topologies required a comparable number of rearrangement steps, making both hypotheses equally likely in terms of chromosomal rearrangements. Similar to previous sequence-based phylogenomic studies (Cunha and Giribet 2019; Uribe et al. 2022; Zhong et al. 2022; Chen et al. 2025b), macrosynteny too have difficulties in resolving the exact branching order of major gastropod lineages. The accumulation of chromosomal rearrangements in the early evolution was likely rapid and complex, potentially obscuring the phylogenetic signal at the macrosyntenic level.

A comparative analysis of chromosomal arrangements across the gastropod subclasses reveals a mosaic of striking genomic stability in some lineages and remarkable flexibility in others. Crucially, a virtually complete spectrum of large-scale genomic rearrangements is demonstrated within Gastropoda, including whole-genome duplication (WGD), chromosome fission, and diverse mechanisms of chromosome fusion, encompassing both end-to-end fusion and fusion-with-mixing. The presence of such a diverse array of structural variations highlights the highly flexible genomes of molluscs (Chen et al. 2025b), and demonstrating it also at the macrosynteny level. In Caenogastropoda, consistent with previous findings in *Pomacea canaliculata* (Xiong et al. 2025), freshwater lineages exhibited extensive end-to-end fusions, whereas deep-sea species remained chromosomally conservative. This contrasts with Heterobranchia, where freshwater species showed conservative genome structures, but terrestrial lineages such as *Arion vulgaris* and *Achatina fulica* have undergone WGD (Guo et al. 2019; Chen et al. 2022), a phenomenon often associated with the physiological demands of terrestrialization (Hallinan and Lindberg 2011). Neritimorpha presents a unique pattern where freshwater colonization is linked to small-scale fusion-with-mixing events and fissions, diverging from the fusion-dominated pattern in Caenogastropoda. The deep-sea vent-endemic vetigastropod *Thermocollonia jamsteci* exhibited extensive fusion-with-mixing events, unlike vent-endemic lineages in other subclasses like Neomphaliones which were characterized by relatively conserved genomes.

Although gastropods have undergone complex chromosomal rearrangements throughout their evolutionary history, these structural variations have not significantly disrupted the ancestral TAD boundaries. Instead, TAD boundaries, acting as critical elements for maintaining higher-order chromatin architecture and the independence of gene regulatory networks, exhibit a high degree of evolutionary conservation. Chromosomal rearrangement events show a strong tendency to treat TADs as intact, independent evolutionary units, with breakpoints, inversions, or translocations predominantly occurring within their interiors, which may change the gene regulation network. This evolutionary paradigm effectively prevents the dysregulation of ectopic interactions among regulatory elements that would otherwise result from boundary disruption, thereby highlighting the profound structural constraints imposed by the three-dimensional genomic architecture on chromosomal evolution.

The functional implications of gastropod chromosomal rearrangements highlight divergent adaptive strategies associated with habitat transitions and colonisations of new environments. Our comparative genomic analysis show that the transition from marine to freshwater ecosystems generally involves far more drastic environmental challenges, such as extreme shifts in osmolarity, than remaining in stable marine habitats including even shifting to deep-sea habitats (Mestre et al. 2008). Similar genomic restructuring driven by severe water-salinity gradients is also evident in plants, which leverage whole-genome duplication and rearrangements for osmoregulation (Wang et al. 2024). In the transition to freshwater from marine habitats, *Pomacea canaliculata* (Caenogastropoda) and *Vittina coromandeliana* (Neritimorpha) adopted fundamentally different approaches. *Pomacea canaliculata* utilized chromosomal rearrangements to integrate adaptive metabolic modules, enhancing energy conversion and detoxification (Xiong et al. 2025). This expansionist strategy likely facilitated its successful transition by conferring the metabolic resilience necessary to overcome physiological barriers. Conversely, *Vittina coromandeliana* employed rearrangements to consolidate homeostatic systems, restructuring immune defense and osmoregulation pathways to ensure physiological stability. This conservative strategy probably reflects prioritizing survival in fluctuating freshwater environments over rapid ecological expansion. In the deep sea, chromosomal rearrangements in *Thermocollonia jamsteci* appears to have helped with multi-layered physiological safeguarding. The enrichment of pathways for mucin biosynthesis and heavy metal detoxification in rearranged regions parallels findings in *Chrysomallon squamiferum* (Sun et al. 2020), reinforcing the hypothesis that genomic plasticity facilitates the evolution of specialized traits required for survival in extreme hydrothermal vent environments.

Our analysis revealed a functional convergence targeting the Mucin type O-glycan biosynthesis pathway in both freshwater and deep-sea hydrothermal vent lineages, yet with distinct adaptations that reflect their specific habitat challenges. While both the freshwater *V. coromandeliana* and the deep-sea Neomphaliones extensively recruited genes encoding the rate-limiting initiation enzymes (GALNT, K00710) and Core 1 synthase (K00731) to establish a fundamental mucosal barrier, *V. coromandeliana* specifically retained additional branching enzymes, including Core 2 (K09662) and Core 3 (K00727) synthases. This increased structural complexity likely enhances the mucin’s capacity for pathogen recognition and immune modulation, critical for the conservative strategy required to maintain homeostasis in freshwater environments. In contrast, Neomphaliones predominantly amplified the initiation modules (K00710), a strategy possibly aimed at maximizing mucin density to form a robust physicochemical shield against heavy metal toxicity and hydrostatic pressure.

In conclusion, we reconstructed the ALGs for Gastropoda and its six subclasses, establishing a robust framework for tracing chromosomal evolution in this extremely diverse animal group. Our results demonstrate that there is no singular evolutionary trajectory of chromosomal rearrangement associated with colonizing new habitat types across Gastropoda. Instead, different lineages have navigated the challenges of the transition to freshwater, terrestrial, and deep-sea environments through fundamentally distinct chromosomal strategies ranging from extensive chromosomal restructuring in some groups including whole-genome duplications to strict conservation in other. The capacity of different gastropod subclasses to adopt independent, diverse, and lineage-specific strategies to adapt to various stressful habitats serve as testaments to the extraordinary genomic flexibility that underlie the immense diversity of Gastropoda.

## Acknowledgments

The authors thank the support of captain and crew of R/V *Falkor (too)* during cruise FKt231024 and R/V *Yokosuka*; and extend the same to the pilots and teams of ROVs *SuBastian*, *KM-ROV*, and *Hercules*. We gratefully acknowledge the chief scientists of the cruises, including John W. Jamieson (University of Ottawa) for FKt231024 and Meghan Paulson (Ocean Networks Canada), Allison Fundis (Ocean Exploration Trust), and Josh Chernov (Ocean Dynamics Inc.). Hidetaka Nomaki (JAMSTEC) is thanked for providing sea-going opportunities on-board R/V *Yokosuka*.

## Data Availability

The raw sequencing reads were deposited in NCBI SRA data base under the BioProject number of PRJNA1519099. The assembled genomes and annotations can be found in Figshare: https://figshare.com/s/2ae63467fec8a87972ea. A graphical representation of spatial preferences of chromosomal rearrangements can be found in Figshare: https://figshare.com/s/8548a51d720f29140923.

