## Supplementary Materials for "Reconstruction of the gastropod ancestral karyotype reveals lineage-specific chromosomal evolution and adaptations during habitat transitions"

The Supplementary Materials file includes:
Figure S1 to S3
Legend for tables S1

Other Supplementary Material for this manuscript includes the following:
Table S1

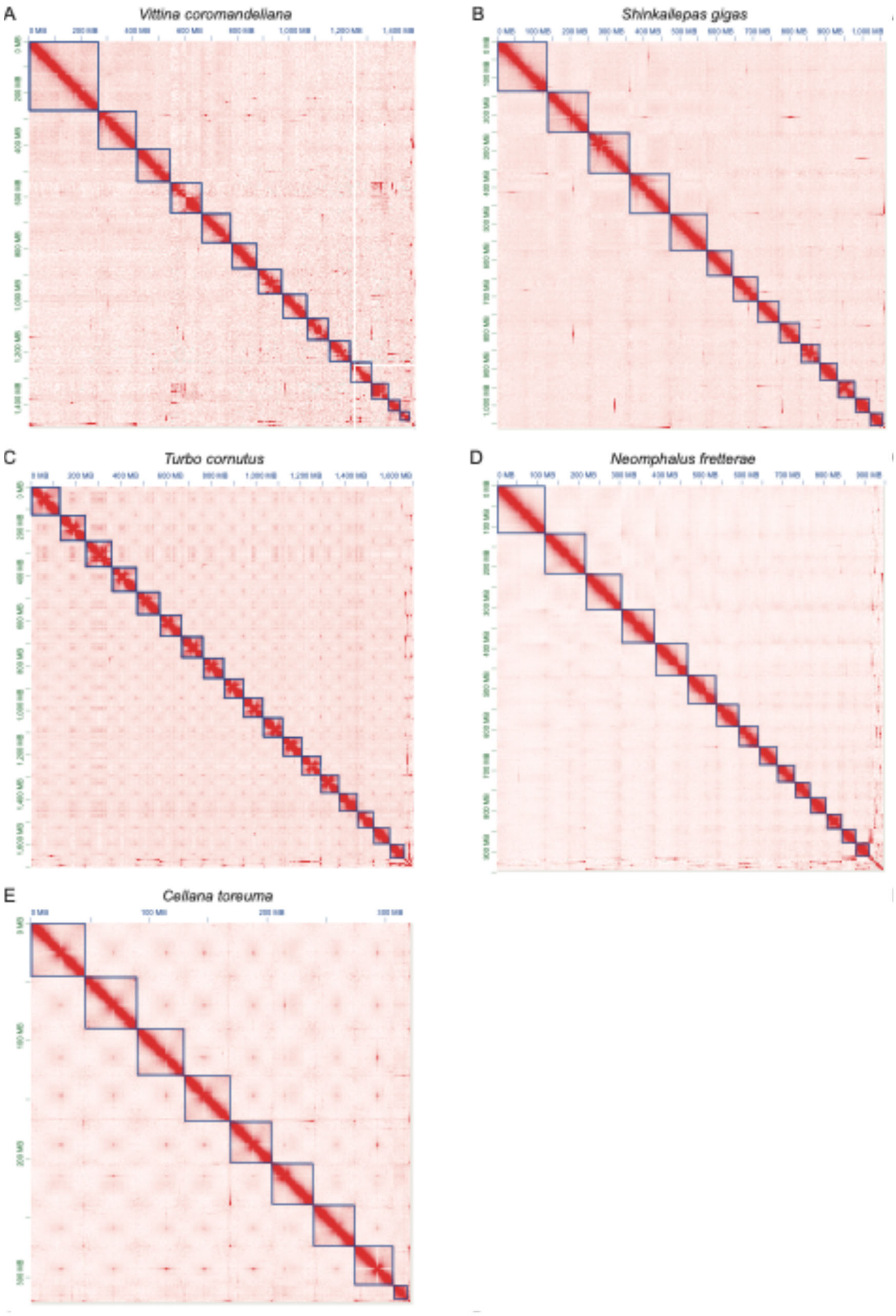


**Figure S1.** Hi-C contact heatmaps for the chromosome-level genome assemblies of the five gastropods for which chromosomal-level genomes were assembled in this study. (A) *Vittina coromandeliana* (Neritimorpha). (B) *Shinkailepas gigas* (Neritimorpha). (C)

*Turbo cornutus* (Vetigastropoda). (D) *Neomphalus fretterae* (Neomphaliones). (E) *Cellana toreuma* (Patellogastropoda).

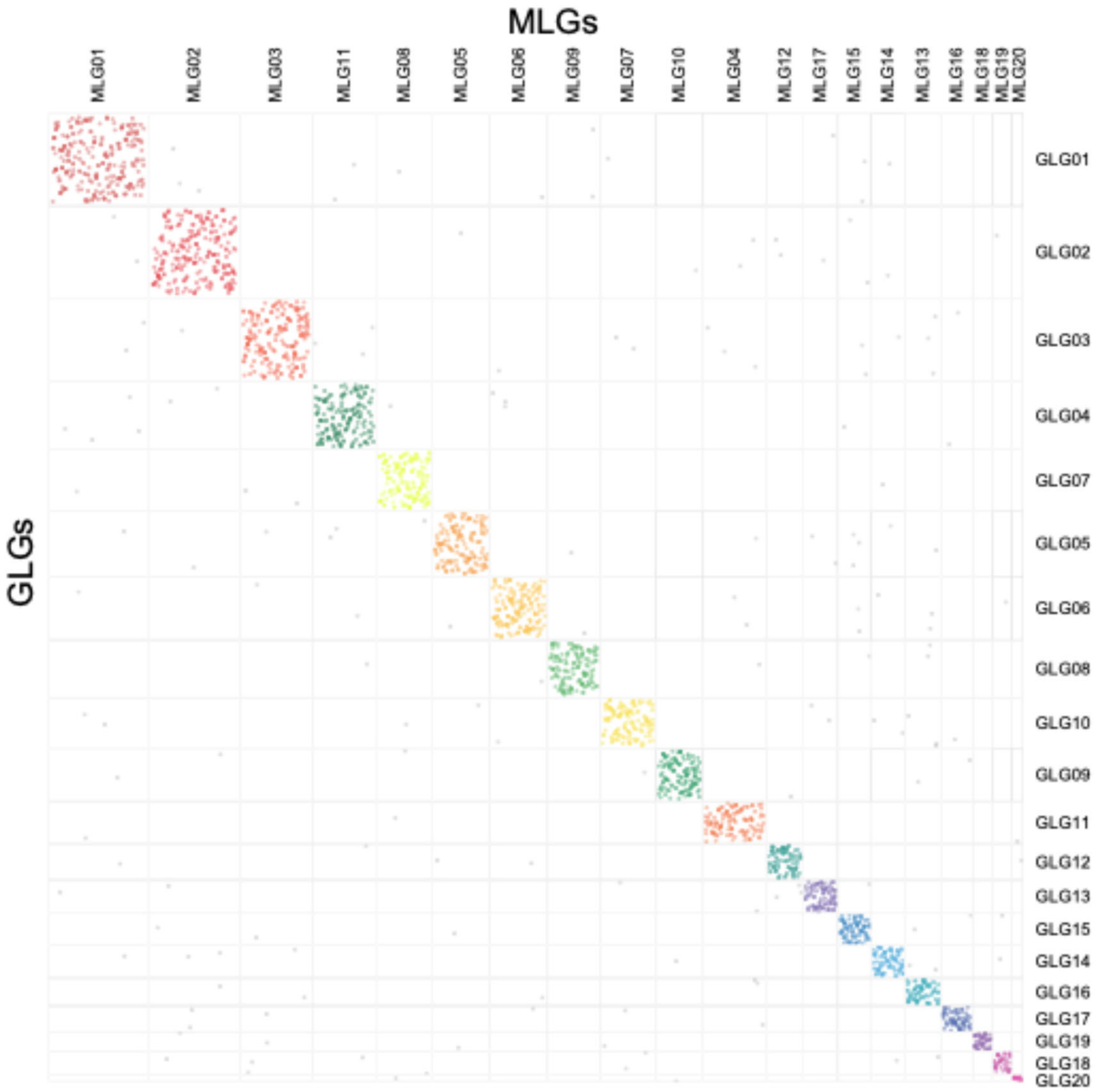


**Figure S2.** Oxford dot-plot of ancestral Gastropod linkage groups (GLGs) against ancestral mollusc linkage groups (MLGs).

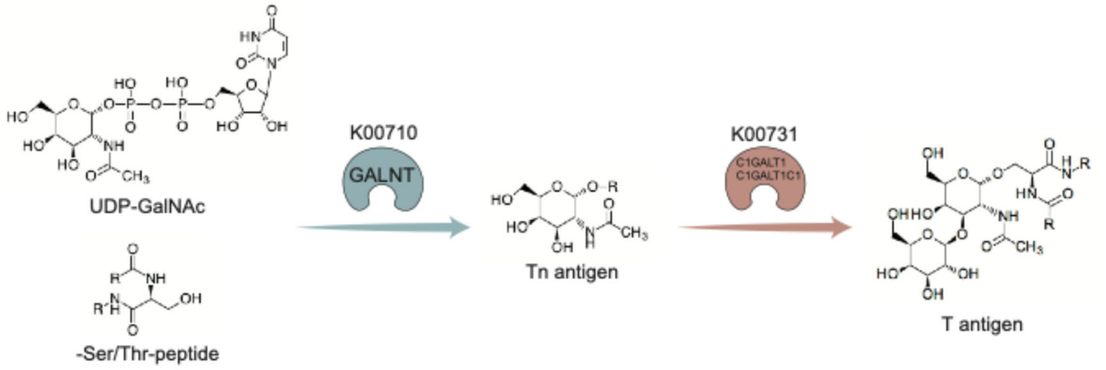


**Figure S3.** Mucin-type O-glycan biosynthesis pathway enriched in rearranged genes in the freshwater neritimorph species *Vittina coromandeliana* and the ancestral linkage groups of Neomphaliones (NpLGs) endemic to the deep sea. This schematic illustrates the core biochemical steps of mucin-type O-glycan synthesis, a pathway significantly enriched within the rearranged genomic regions of the freshwater species *Vittina coromandeliana* and NpLGs. The process initiates with the transfer of GalNAc from UDP-GalNAc to Ser/Thr residues catalyzed by the polypeptide N-acetylgalactosaminyltransferases (GALNTs, KO: K00710), forming the Tn antigen; subsequently, the Tn antigen is elongated into the T antigen by the action of C1GALT1 and its chaperone C1GALT1C1 (KO: K00731).

**Legend for tables S1**

**Table S1.** Summary of genomes used in this study.
